# High-quality reference genome for an arid-adapted mammal, the banner-tailed kangaroo rat (*Dipodomys spectabilis*)

**DOI:** 10.1101/2021.11.23.469702

**Authors:** Avril M. Harder, Kimberly K.O. Walden, Nicholas J. Marra, Janna R. Willoughby

## Abstract

Kangaroo rats in the genus *Dipodomys* are found in a variety of habitat types in western North America, including deserts, arid and semi-arid grasslands, and scrublands. Many *Dipodomys* species are experiencing strong population declines due to increasing habitat fragmentation, with two species listed as federally endangered. The precarious state of many *Dipodomys* populations, including those occupying extreme environments, make species of this genus valuable subjects for studying the impacts of habitat degradation and fragmentation on population genomic patterns and for characterizing the genomic bases of adaptation to harsh conditions. To facilitate exploration of such questions, we assembled and annotated a reference genome for the banner-tailed kangaroo rat (*D. spectabilis*) using PacBio HiFi sequencing reads, providing a more contiguous genomic resource than two previously assembled *Dipodomys* genomes. Using the HiFi data for *D. spectabilis* and publicly available sequencing data for two other *Dipodomys* species (*D. ordii* and *D. stephensi*), we demonstrate the utility of this new assembly for studies of congeners by conducting inference of historic effective population sizes (*N*_e_) and linking these patterns to the species’ current extinction risk statuses. The genome assembly presented here will serve as a valuable resource for population and conservation genomic studies of *Dipodomys* species, comparative genomic research within mammals and rodents, and investigations into genomic adaptation to extreme environments and changing landscapes.

**Significance statement:** Kangaroo rats in the genus *Dipodomys* occur in a wide variety of habitat types, ranging from scrublands to arid deserts, and are increasingly impacted by habitat fragmentation with populations of many species in strong decline. To facilitate population and conservation genomic studies of *Dipodomys* species, we generated the first reference genome assembly for the extensively studied banner-tailed kangaroo rat (*D. spectabilis*) from long read PacBio sequencing data. The genome assembly presented here will serve as a valuable resource for studies of *Dipodomys* species—which have long served as ecological and physiological models for the study of osmoregulation—comparative genomic surveys of mammals and rodents, and investigations into genomic adaptation to extreme environments and changing landscapes.

## Introduction

The genus *Dipodomys* comprises 20 species of kangaroo rats and belongs to the Heteromyidae family of rodents (Alexander & Riddle 2005). Often considered keystone species, kangaroo rats inhabit warm and cold deserts, arid and semi-arid grasslands, and scrublands of western North America, with most species preferring sandy soils that allow for construction of elaborate underground burrows used for protection from predators and harsh environmental conditions, reproduction, and food caching (Alexander & Riddle 2005; Brown & Heske 1990). For many of these species, limited dispersal capabilities and habitat fragmentation has led to population declines, with five species classified as threatened by IUCN (International Union for Conservation of Nature) and two species (and four additional subspecies) listed as federally endangered, of which several have been the subject of management efforts to decrease species vulnerability (IUCN 2021; Shier et al. 2021; Hendricks et al. 2020; Blackhawk et al. 2016; Loew et al. 2006). The imperiled state of many *Dipodomys* populations, combined with the vast array of occupied habitat types that include arid environments, make species of this genus valuable subjects for studying the impacts of habitat degradation and fragmentation on population genomic patterns and for characterizing the genomic bases of adaptation to extreme environments. Specifically, the latter application may provide a key link between historical studies of *Dipodomys* kidney morphology (Vimtrup & Schmidt-Nielsen 1952; Schmidt-Nielsen & Schmidt-Nielsen 1952) and more recent investigations of gene and protein expression related to osmoregulation in arid conditions (reviewed in Rocha et al. 2021).

To facilitate genomic studies of *Dipodomys* species, we used PacBio HiFi sequencing data to generate an annotated reference genome for the banner-tailed kangaroo rat (*D. spectabilis*), a species whose ecology and evolution has been studied extensively for several decades (Vorhies & Taylor 1922; Greene & Reynard 1932; Schroder 1979; Waser & Jones 1991; Busch et al. 2007; Willoughby et al. 2019). Prior to the assembly of this genome, two *Dipodomys* reference genomes were available that had been built with short read sequencing data, with one assembly comprising 1.3 million scaffolds totaling 2.3 Gb (*D. stephensi*, GCA_004024685.1) and the other comprising 65,193 scaffolds totaling 2.2 Gb (*D. ordii*, GCA_000151885.2). We used long read sequencing data to generate a *D. spectabilis* assembly with increased contiguity and then used short read sequences from *D. ordii* and *D. stephensi* to conduct historical effective population size (*N*_e_) inference for all three species. We compared population trajectories for these two species and *D. spectabilis* to identify demographic trends associated with current extinction risk designations of the species, which range from IUCN listings of least concern (*D. ordii*) and near threatened (*D. spectabilis*) to vulnerable and U.S. federally endangered (*D. stephensi*). These analyses highlight the utility of the *D. spectabilis* reference genome for exploring evolutionary questions with conservation implications.

## Results and Discussion

### Genome assembly and annotation

We generated 97.9 Gb of PacBio HiFi reads resulting in 23X coverage of the final assembly. The hifiasm primary contig assembly summed to 2.8 Gb and comprised 2,026 contigs with an assembly contig N50 of 9.6 Mb (Table 1). Prior to annotation, Benchmarking Universal Single-Copy Orthologs (BUSCO) v5.1.2 assessment identified 97.2% of vertebrata_odb10 orthologs as complete (94.2% single-copy, 3.0% duplicated), 1.1% as fragmented, and 1.7% as missing.Identification of repetitive sequences with WindowMasker resulted in the masking of 47.85% of the genome sequence. Annotation of the masked genome via NCBI’s pipeline identified 20,632 protein-coding genes with a mean of 1.66 transcripts per gene and an average gene length of 31,775 bp. The BUSCO v4.1.4 results for the annotated gene set classified 98.2% of glires_odb10 orthologs as complete (96.5% single-copy, 1.7% duplicated), 0.4% as fragmented, and 1.5% as missing.

**Table 1.**
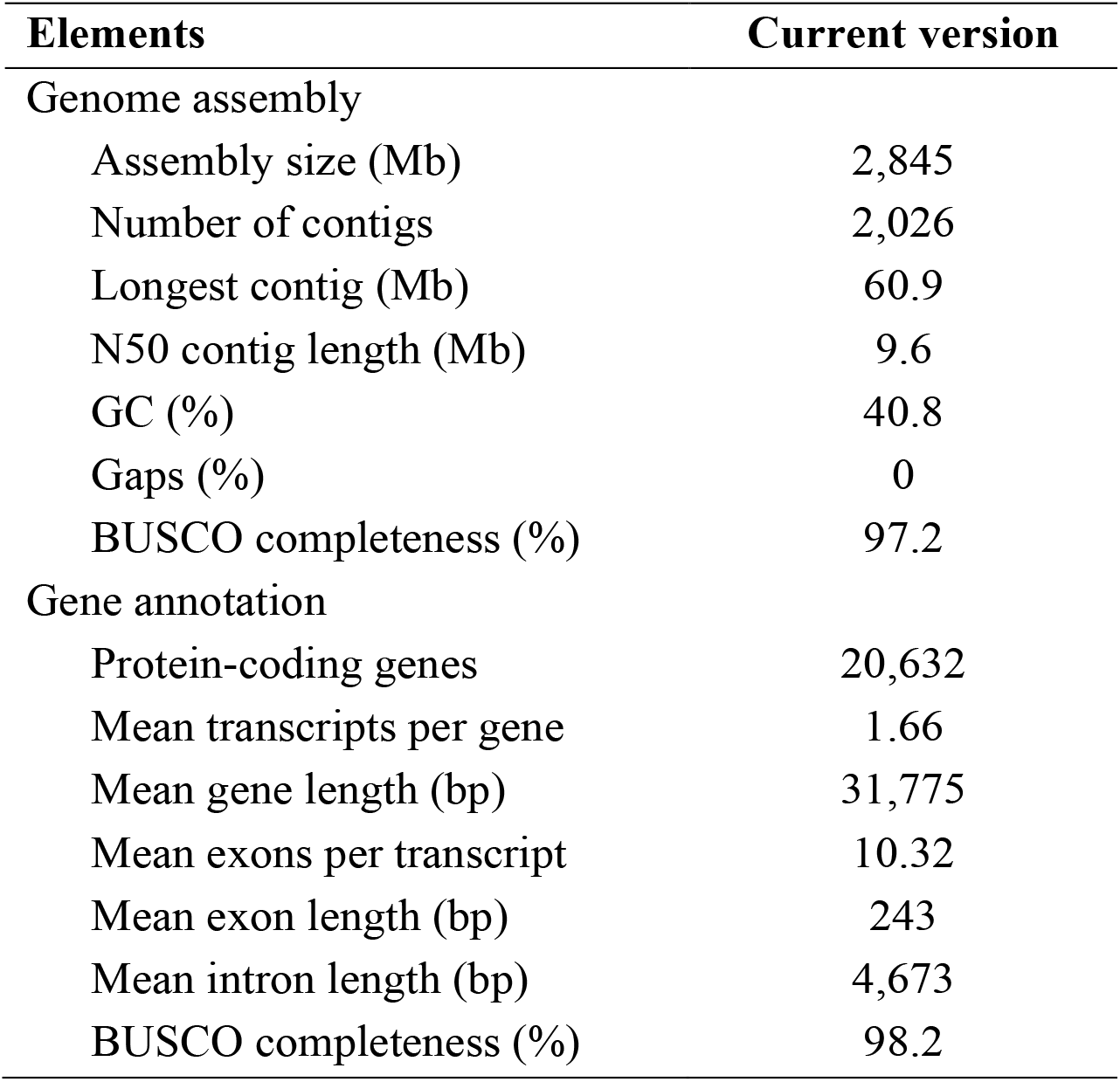
Genome assembly and annotation statistics for *Dipodomys spectabilis*.

| Elements | Current version |
| --- | --- |
| Genome assembly |  |
| Assembly size (Mb) | 2,845 |
| Number of contigs | 2,026 |
| Longest contig (Mb) | 60.9 |
| N50 contig length (Mb) | 9.6 |
| GC (%) | 40.8 |
| Gaps (%) | 0 |
| BUSCO completeness (%) | 97.2 |
| Gene annotation |  |
| Protein-coding genes | 20,632 |
| Mean transcripts per gene | 1.66 |
| Mean gene length (bp) | 31,775 |
| Mean exons per transcript | 10.32 |
| Mean exon length (bp) | 243 |
| Mean intron length (bp) | 4,673 |
| BUSCO completeness (%) | 98.2 |

Prior to assembly of the *D. spectabilis* genome, the most contiguous *Dipodomys* reference genome was built for *D. ordii* (GCA_000151885.2). The *D. spectabilis* assembly is nearly 800 Mb larger than the ungapped *D. ordii* assembly with a markedly improved contig N50 (9.6 Mb) over *D. ordii* (48,087 bp) and is organized into far fewer sequences (2,026 contigs) than the *D. ordii* assembly (148,226 contigs organized into 65,193 scaffolds). Furthermore, the longest contigs in the *D. spectabilis* assembly (maximum = 60.9 Mb) approach lengths typical of mammalian chromosome arms (Klegarth & Eisenberg 2018). The new *D. spectabilis* annotation also comprises a greater number of fully-supported mRNA sequences than the *D. ordii* assembly (33,775 vs. 22,964) and will provide increased resolution to genomic studies of *Dipodomys* that aim to describe patterns of adaptation and signals of selection.

### Variant calling

For both *D. ordii* and *D. stephensi*, read mapping rates to the *D. spectabilis* reference genome were high (95.9% and 92.1%, respectively), demonstrating the utility of this assembly for genomic studies across the genus despite the approximately 10 million years separating these three species from their most recent common ancestor (Figure 1A). After the final filtering step for PSMC (*i*.*e*., retaining sites with read depths greater than 1/3 and less than two times the average depth of coverage within a sample), we retained genotypes for a total of 2.3 billion sites for *D. spectabilis*, 1.7 billion for *D. stephensi*, and 1.7 billion for *D. ordii*. The proportion of heterozygous sites was lowest for *D. ordii* (0.0012) and was similar between *D. spectabilis* (0.0021) and *D. stephensi* (0.0023). Mapping to a congeneric reference genome could result in biased estimates of heterozygosity. Because *D. ordii* and *D. stephensi* are more closely related to one another than either species is to *D. spectabilis* (Figure 1A), mapping bias is expected to affect estimates for *D. ordii* and *D. stephensi* in the same direction (i.e., either under- or overestimation) and with similar magnitude. However, our heterozygosity results are not consistent with such a pattern. For *D. ordii*, the heterozygosity result contrasts with the much larger range documented for *D. ordii* than for the other two species (Figure 1B), but the low heterozygosity of the *D. ordii* sample likely reflects the recent demographic history of the sequenced individual’s population of origin, rather than long-term, species-level trends.

**Figure 1.**
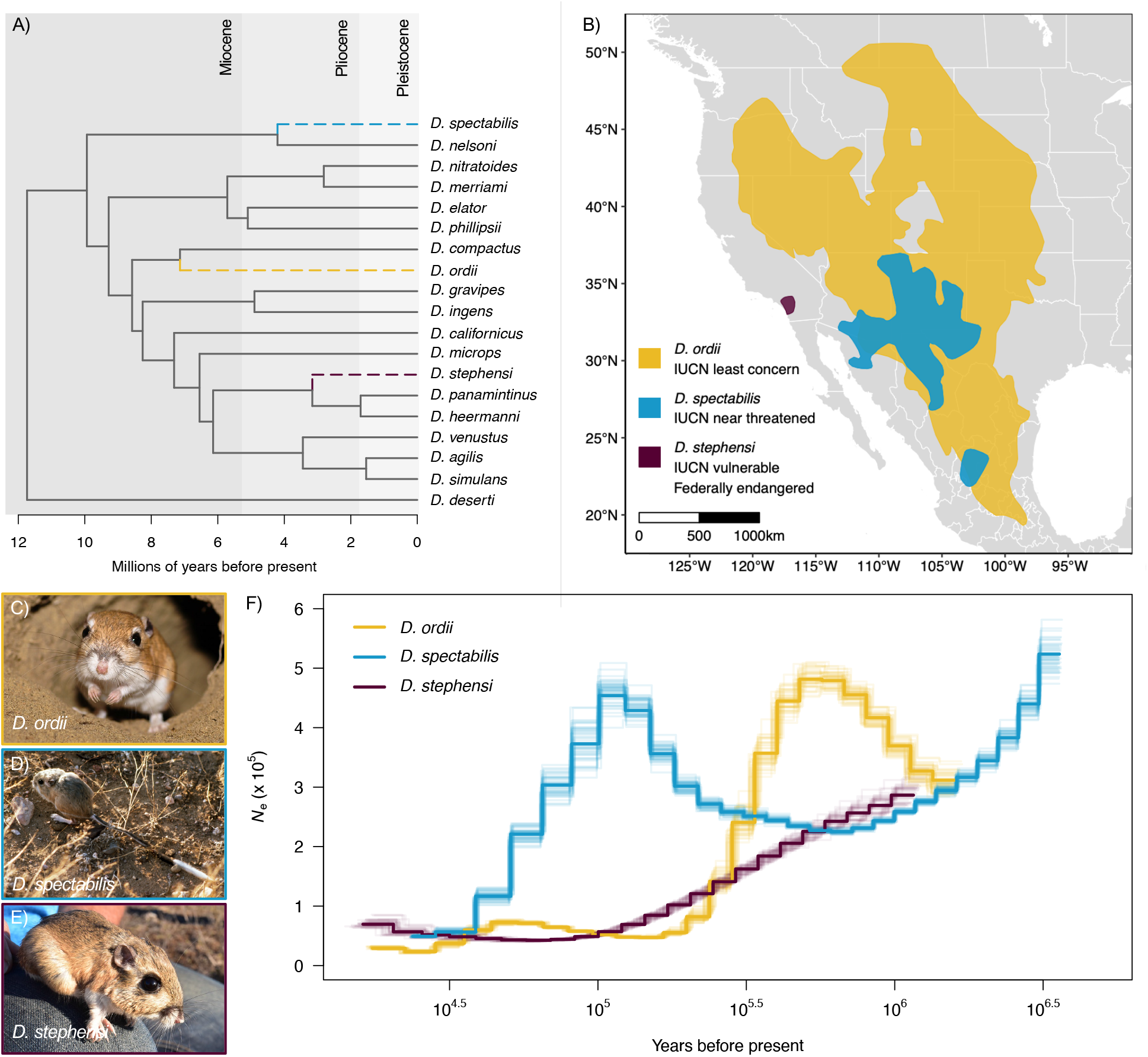
A) *Dipodomys* phylogeny constructed using the TimeTree database. The most recent common ancestor of *D. spectabilis, D. ordii*, and *D. stephensi* lived approximately 10 million years ago. B) Current ranges and conservation statuses for *D*. ordii, *D. spectabilis*, and *D. stephensi* in western North America. C-E) Photos of *D. ordii, D. spectabilis*, and *D. stephensi*. F) PSMC results for all three species, plotted using a generation time of 1 year and a substitution rate of 2.2e-9 per base pair per year. Lighter lines represent results of 50 bootstrap replicates per species. Map made with Natural Earth and the *rnaturalearth* package in R using the WGS84 coordinate system. Species range shapefiles downloaded from IUCN (IUCN 2008, 2009, 2018). Image credits: Andy Teucher (*D. ordii*; https://bit.ly/3Cpcjfl), Peter Waser (*D. spectabilis*), USFWS Pacific Southwest Region (*D. stephensi*;https://bit.ly/3nsHPVu).

### Historical effective population size inference

Each of the three species we analyzed displays distinct patterns of historical *N*_e_ as inferred from PSMC analysis (Figure 1F). *D. ordii* and *D. spectabilis* both exhibit peaks in historical *N*_e_ followed by declines, whereas *D. stephensi* appears to have experienced steadily declining *N*_e_ over the time range covered by our analysis. The range of time over which changes in *N*_e_ were inferred for *D. spectabilis* extends approximately 2 million years farther into the past than the time ranges for *D. ordii* and *D. stephensi*. The increased temporal range of PSMC results for *D. spectabilis* relative to the other two species is likely due to the use of long-read data to identify variants, which allows for resolution of longer haplotypes and, consequently, inference of older coalescent events compared to when variants are called from short-read data.

Initially, diversification within heteromyid rodents likely took place towards the middle of the Miocene (Figure 1A), around the time that habitat types were also diversifying across their western North America ranges. Further speciation within *Dipodomys* coincided with continuing partitioning of ecoregions and cyclical formation of glacial refugia into the Pleistocene (Alexander & Riddle 2005). Although all three species exhibit declines in historical *N*_e_, the steady decline observed for *D. stephensi* is consistent with this species’ extremely limited current range and U.S. federally endangered status. Both *D. spectabilis* and *D. ordii* maintain much larger contemporary range sizes than *D. stephensi* and are of relatively less conservation concern. Declines in *N*_e_ for these species may correspond to disrupted movement patterns among populations during Pleistocene glacial-interglacial cycles.

Through these analyses, we have demonstrated the applicability of our new *D. spectabilis* reference genome to genomic studies across the genus *Dipodomys*, a taxon comprising species with a broad range of conservation statuses and habitat preferences. Our annotated assembly represents a significant improvement in resources for this genus and will facilitate future investigations into a broad range of eco-evolutionary questions, including for species of conservation concern, such as *D. stephensi*, or for other *Dipodomys* species experiencing increasing habitat fragmentation and population declines (Blackhawk et al. 2016; Hendricks et al. 2020).

## Materials and Methods

### Sample collection and sequencing

The male *Dipodomys spectabilis* individual used for genomic sequencing was collected near Portal, AZ (31°56’N, 109°5’W) in December 2009. Whole organs (kidney and liver) were dissected and frozen in liquid nitrogen for transport to Purdue University and subsequent storage at -80 °C (Marra et al. 2014, 2012). In February 2021, 0.1 g portions of both tissues were shipped on dry ice to Polar Genomics (Ithaca, NY) for extraction of high-molecular weight genomic DNA (gDNA) using a modified nuclei extraction protocol (Zhang et al. 2012). Extracted gDNA was supplied to the Roy J. Carver Biotechnology Center DNA services core (University of Illinois) where it was sheared to an average fragment length of 13kb with a Megaruptor 3.

Sheared gDNA was converted to a PacBio library with the SMRTBell Express Template Prep Kit 2.0, and this library was sequenced on 6 SMRT cells 8M on a PacBio Sequel IIe using the circular consensus sequencing (CCS) mode and a 30-hour movie time to produce HiFi reads. CCS analysis was performed using SMRTLink V10.0 using default parameters.

### Genome assembly and annotation

PacBio HiFi reads were filtered to retain those between 4 kb and 40 kb with seqkit (Shen et al. 2016) and then assembled with hifiasm v.0.14.2 (r315) using default parameters (Cheng et al. 2021). Genome completeness was assessed using BUSCO v.5.1.2 in “genome” mode with the vertebrata_odb10 lineage dataset (3,354 orthologs) (Manni et al. 2021).

Annotation of genes and genomic features from the hifiasm primary contigs was conducted by NCBI via the Eukaryotic Genome Annotation Pipeline v9.0 (release date June 8 2021) (Thibaud-Nissen et al. 2013). This pipeline includes repeat masking using WindowMasker (Morgulis et al. 2006) and gene model prediction for the masked genome informed by NCBI RefSeq transcript and protein sets for *Mus musculus* and publicly available RNA-Seq data. The RNA-Seq reads used for gene prediction originated from kidney and spleen tissues of *D. spectabilis* and two other heteromyid rodents (*Heteromys desmarestianus* and *Chaetodipus baileyi*). The details of the NCBI annotation pipeline can be found at: https://www.ncbi.nlm.nih.gov/genome/annotation_euk/process/. The final annotation (NCBI *Dipodomys spectabilis* Annotation Release 100) was assessed using BUSCO v4.1.4 in “protein” mode using the glires_odb10 lineage data set (13,798 orthologs) (Simão et al. 2015).

### Variant calling

We downloaded Illumina sequencing reads from NCBI’s Sequence Read Archive (SRA) that were previously used in assembly of the *D. ordii* and *D. stephensi* reference genomes (*D. ordii* sequence accessions SRR1646412-23 and *D. stephensi* sequence accession SRR14572526). We removed adapter sequences and clipped low quality bases (quality score < 20) from both ends of reads using Trimmomatic v0.39 (Bolger et al. 2014). Reads were quality-checked before and after trimming using FastQC v0.11.9 (Andrews 2015) and were aligned to the newly assembled *D. spectabilis* reference genome using the BWA-MEM algorithm implemented in BWA v0.7.12 (Li 2013). To align our HiFi reads for the *D. spectabilis* individual to the assembled genome, we used the map-hifi option in Minimap2 (Li 2018). When necessary, we merged resulting BAM files into a single file per species using SAMtools v1.11 (Li et al. 2009). Finally, we sorted the BAM files and calculated average depth of coverage across each contig, again using SAMtools.

For each species, a variant file was produced using the BCFtools v1.10.2 mpileup and call commands (Danecek et al. 2021). We initially filtered each VCF file to exclude indels (*i*.*e*., retain SNPs) and to retain only sites with quality scores > 30 and minimum read depth of 15 using VCFtools v0.1.14 (Danecek et al. 2011). Variants were further limited to those located on contigs ≥ 100 kb in length and contigs likely not originating from sex chromosomes, leaving variants on 652 contigs totaling 2.55 Gb in length (89.7% of reference assembly).

### Inference of historical effective population sizes

To compare patterns of historical effective population size (*N*_e_) among *Dipodomys* species, we began by using the BCFtools vcfutils.pl script and the PSMC fq2psmcfa function (Li & Durbin 2011) to convert the VCF files to masked consensus FASTA files. For each file, we applied a final variant filter to exclude sites with read depths less than 1/3 and greater than two times the average depth of coverage calculated for each sample above, as recommended by the PSMC authors (https://github.com/lh3/psmc), and determined the proportion of heterozygous sites.

After first testing default PSMC parameter settings, we ultimately set the -p parameter to ‘8+25^*^2+2+4’ and confirmed that at least ten recombination events were inferred in each interval within 20 iterations before also performing 50 iterations of bootstrapping. We plotted the results in R v4.0.3 (R Core Team 2020), supplying the average mammalian mutation rate (2.2e-9 per base pair per year) (Kumar & Subramanian 2002) and a generation time of 1 year to the functions defined in the plotPsmc.R script published on Dryad (Liu & Hansen 2017, 2016). To place the PSMC results into broader taxonomic context, we also constructed a *Dipodomys* phylogeny using a Newick file exported from the TimeTree database (Kumar et al. 2017) and plotted using the *ape* package in R.

## Data Availability

Genome assembly and raw sequencing data have been deposited at the NCBI under the accessions GCA_019054845.1 and SRX11001182-SRX11001186, respectively. Genome annotation is available under the NCBI accession ASM1905484v1. Code for all bioinformatic analyses available at https://github.com/avril-m-harder/D_spectabilis_genome_resource and https://github.com/jwillou/ D_spectabilis_genome_resource.

## Acknowledgements

This material is based on work supported by the National Science Foundation Postdoctoral Research Fellowships in Biology Program under Grant No. 2010251 to AMH. This work was also supported by the U.S. Department of Agriculture, National Institute of Food and Agriculture, Hatch project 1025651 to JRW. We thank J. Vrebalov of Polar Genomics for performing DNA extraction and the University of Illinois’ Roy J. Carver Biotechnology Center DNA Services Lab for library construction and sequencing.

## Notes

https://github.com/avril-m-harder/D_spectabilis_genome_resource

### Competing Interest Statement

The authors have declared no competing interest.

## Literature Cited

Alexander LF, Riddle BR. 2005. Phylogenetics of the New World rodent family Heteromyidae. Journal of Mammalogy. 86:366–379. doi: 10.1644/BER-120.1.

Andrews S. 2015. FastQC: A quality control tool for high throughput sequence data. http://www.bioinformatics.babraham.ac.uk/projects/fastqc/.

Blackhawk NC, Germano DJ, Smith PT. 2016. Genetic variation among populations of the endangered giant kangaroo rat, Dipodomys ingens, in the southern San Joaquin Valley. The American Midland Naturalist. 175:261–274. doi: 10.1674/0003-0031-175.2.261.

Bolger AM, Lohse M, Usadel B. 2014. Trimmomatic: a flexible trimmer for Illumina sequence data. Bioinformatics. 30:2114–2120. doi: 10.1093/bioinformatics/btu170.

Brown JH, Heske EJ. 1990. Control of a desert-grassland transition by a keystone rodent guild. Science. 250:1705–1707. doi: 10.1126/science.250.4988.1705.

Busch JD, Waser PM, DeWoody JA. 2007. Recent demographic bottlenecks are not accompanied by a genetic signature in banner-tailed kangaroo rats (Dipodomys spectabilis). Molecular Ecology. 16:2450–2462.

Cheng H, Concepcion GT, Feng X, Zhang H, Li H. 2021. Haplotype-resolved de novo assembly using phased assembly graphs with hifiasm. Nature Methods. 18:170–175.

Danecek P et al. 2011. The variant call format and VCFtools. Bioinformatics. 27:2156–2158. doi: 10.1093/bioinformatics/btr330.

Danecek P et al. 2021. Twelve years of SAMtools and BCFtools. GigaScience. 10:giab008. doi: 10.1093/gigascience/giab008.

Greene RA, Reynard C. 1932. The influence of two burrowing rodents, Dipodomys spectabilis spectabilis (kangaroo rat) and Neotoma albigula albigula (pack rat) on desert soils in Arizona. Ecology. 13:73–80. doi: 10.2307/1932493.

Hendricks S et al. 2020. Patterns of genetic partitioning and gene flow in the endangered San Bernardino kangaroo rat (Dipodomys merriami parvus) and implications for conservation management. Conservation Genetics. 21:819–833. doi: 10.1007/s10592-020-01289-z.

IUCN 2008. Dipodomys ordii. The IUCN Red List of Threatened Species. Version 2021-2

IUCN SSC Small Mammal Specialist Group 2018. Dipodomys stephensi. The IUCN Red List of Threatened Species. Version 2021-2

IUCN SSC Small Mammal Specialist Group 2019. Dipodomys spectabilis. The IUCN Red List of Threatened Species. Version 2021-2

IUCN 2021. The IUCN Red List of Threatened Species. https://www.iucnredlist.org (Accessed November 1, 2021).

Klegarth AR, Eisenberg DTA. 2018. Mammalian chromosome–telomere length dynamics. Royal Society Open Science. 5:180492. doi: 10.1098/rsos.180492.

Kumar S, Stecher G, Suleski M, Hedges SB. 2017. TimeTree: A resource for timelines, timetrees, and divergence times. Molecular Biology and Evolution. 34:1812–1819. doi: 10.1093/molbev/msx116.

Kumar S, Subramanian S. 2002. Mutation rates in mammalian genomes. Proceedings of the National Academy of Sciences. 99:803–808. doi: 10.1073/pnas.022629899.

Li H. 2013. Aligning sequence reads, clone sequences and assembly contigs with BWA-MEM. 1303.3997 [q-bio]. http://arxiv.org/abs/1303.3997 (Accessed May 8, 2017).

Li H. 2018. Minimap2: pairwise alignment for nucleotide sequences. Bioinformatics. 34:3094–3100. doi: 10.1093/bioinformatics/bty191.

Li H et al. 2009. The sequence alignment/map format and SAMtools. Bioinformatics. 25:2078–2079. doi: 10.1093/bioinformatics/btp352.

Li H, Durbin R. 2011. Inference of human population history from individual whole-genome sequences. Nature. 475:493–496. doi: 10.1038/nature10231.

[dataset] * Liu S, Hansen MM. 2016. Data from: PSMC (pairwise sequentially Markovian coalescent) analysis of RAD (restriction site associated DNA) sequencing data. Dryad. Dataset. doi: https://doi.org/10.5061/dryad.0618v.

Liu S, Hansen MM. 2017. PSMC (pairwise sequentially Markovian coalescent) analysis of RAD (restriction site associated DNA) sequencing data. Molecular Ecology Resources. 17:631–641. doi: 10.1111/1755-0998.12606.

Loew SS, Williams DF, Ralls K, Pilgrim K, Fleischer RC. 2006. Population structure and genetic variation in the endangered Giant Kangaroo Rat (Dipodomys ingens). Conservation Genetics. 6:495–510. doi: 10.1007/s10592-005-9005-9.

Manni M, Berkeley MR, Seppey M, Simão FA, Zdobnov EM. 2021. BUSCO update: novel and streamlined workflows along with broader and deeper phylogenetic coverage for scoring of eukaryotic, prokaryotic, and viral genomes. 2106.11799.

Marra NJ, Eo SH, Hale MC, Waser PM, DeWoody JA. 2012. A priori and a posteriori approaches for finding genes of evolutionary interest in non-model species: Osmoregulatory genes in the kidney transcriptome of the desert rodent Dipodomys spectabilis (banner-tailed kangaroo rat). Comparative Biochemistry and Physiology Part D: Genomics and Proteomics. 7:328–339. doi: 10.1016/j.cbd.2012.07.001.

Marra NJ, Romero A, DeWoody JA. 2014. Natural selection and the genetic basis of osmoregulation in heteromyid rodents as revealed by RNA-seq. Molecular Ecology. 23:2699–2711. doi: 10.1111/mec.12764.

Morgulis A, Gertz EM, Schaffer AA, Agarwala R. 2006. WindowMasker: window-based masker for sequenced genomes. Bioinformatics. 22:134–141. doi: 10.1093/bioinformatics/bti774.

R Core Team. 2020. R: a language and environment for statistical computing. R Foundation for Statistical Computing: Vienna, Austria https://www.R-project.org/.

Rocha JL, Godinho R, Brito JC, Nielsen R. 2021. Life in deserts: The genetic basis of mammalian desert adaptation. Trends in Ecology & Evolution. 36:637–650. doi: 10.1016/j.tree.2021.03.007.

Schmidt-Nielsen K, Schmidt-Nielsen B. 1952. Water metabolism of desert mammals. Physiological Reviews. 32:135–166. doi: 10.1152/physrev.1952.32.2.135.

Schroder GD. 1979. Foraging behavior and home range utilization of the bannertail kangaroo rat (Dipodomys spectabilis). Ecology. 60:657–665. doi: 10.2307/1936601.

Shen W, Le S, Li Y, Hu F. 2016. SeqKit: A cross-platform and ultrafast toolkit for FASTA/Q file manipulation. PLoS ONE. 11:e0163962. doi: 10.1371/journal.pone.0163962.

Shier DM et al. 2021. Genetic and ecological evidence of long term translocation success of the federally endangered Stephens’ kangaroo rat. Conservation Science and Practice. 3. doi: 10.1111/csp2.478.

Simão FA, Waterhouse RM, Ioannidis P, Kriventseva EV, Zdobnov EM. 2015. BUSCO: assessing genome assembly and annotation completeness with single-copy orthologs. Bioinformatics. 31:3210–3212. doi: 10.1093/bioinformatics/btv351.

Thibaud-Nissen F, Souvorov A, Murphy T, DiCuccio M, Kitts P. 2013. Eukaryotic Genome Annotation Pipeline. In: The NCBI Handbook [Internet]. Bethesda (MD): National Center for Biotechnology Information.

Vimtrup BJ, Schmidt-Nielsen B. 1952. The histology of the kidney of kangaroo rats. The Anatomical Record 114:515–528. doi: 10.1002/ar.1091140402.

Vorhies CT, Taylor WP. 1922. Life history of the kangaroo rat, Dipodomys spectabilis spectabilis Merriam. U.S. Department of Agriculture Bulletin. 1901:1–40.

Waser PM, Jones WT. 1991. Survival and reproductive effort in banner-tailed kangaroo rats. Ecology. 72:771–777. doi: 10.2307/1940579.

Willoughby JR, Waser PM, Brüniche-Olsen A, Christie MR. 2019. Inbreeding load and inbreeding depression estimated from lifetime reproductive success in a small, dispersal-limited population. Heredity. 123:192–201. doi: 10.1038/s41437-019-0197-z.

Zhang M et al. 2012. Preparation of megabase-sized DNA from a variety of organisms using the nuclei method for advanced genomics research. Nature Protocols. 7:467–478. doi: 10.1038/nprot.2011.455.

